# *In vitro* evolution of Remdesivir resistance reveals genome plasticity of SARS-CoV-2

**DOI:** 10.1101/2021.02.01.429199

**Authors:** Agnieszka M. Szemiel, Andres Merits, Richard J. Orton, Oscar MacLean, Rute Maria Pinto, Arthur Wickenhagen, Gauthier Lieber, Matthew L. Turnbull, Sainan Wang, Daniel Mair, Ana da Silva Filipe, Brian J. Willett, Sam J. Wilson, Arvind H. Patel, Emma C. Thomson, Massimo Palmarini, Alain Kohl, Meredith E. Stewart

## Abstract

Remdesivir (RDV) is used widely for COVID-19 patients despite varying results in recent clinical trials. Here, we show how serially passaging SARS-CoV-2 *in vitro* in the presence of RDV selected for drug-resistant viral populations. We determined that the E802D mutation in the RNA-dependent RNA polymerase was sufficient to confer decreased RDV sensitivity without affecting viral fitness. Analysis of more than 200,000 sequences of globally circulating SARS-CoV-2 variants show no evidence of widespread transmission of RDV-resistant mutants. Surprisingly, we also observed changes in the Spike (i.e., H69 E484, N501, H655) corresponding to mutations identified in emerging SARS-CoV-2 variants indicating that they can arise *in vitro* in the absence of immune selection. This study illustrates SARS-CoV-2 genome plasticity and offers new perspectives on surveillance of viral variants.

**One Sentence Summary:** SARS-CoV-2 drug resistance & genome plasticity

## Main Text

The Covid-19 pandemic has caused more than 2 million deaths and placed the global economy under considerable strain *(1*). The global effort to repurpose antiviral inhibitors and antiinflammatory compounds to stem virus replication and clinical pathology identified Remdesivir (RDV), a broadly acting nucleoside analogue, as a frontline treatment for patients hospitalized with severe acute respiratory syndrome virus-2 (SARS-CoV-2). RDV exhibits a potent ability to restrict virus replication *in vitro* (*2, 3*). Three randomized trials (*4–6*) demonstrated that RDV treatment reduced recovery time by 31% and demonstrated a non-significant trend towards lower mortality, thus reducing long-term healthcare costs. This trend of reduced hospitalization time and decreased morbidity was further supported by smaller non-randomized studies (*7*). Conversely, a larger trial conducted by WHO (Solidarity Therapeutics Trial) reported no effect on patient survival (*8*). The timing of administration of RDV appeared to be critical for its efficacy (*3, 9, 10*). Despite these inconsistent findings, countries including the USA and UK routinely use RDV for the treatment of hospitalized SARS-CoV-2 patients requiring oxygen who are still within the virological phase of infection (<10 days of illness). RDV is often prescribed in combination with dexamethasone, a steroid treatment, which reduces mortality in ventilated patients (*11, 12*). However, RDV and dexamethasone have yet to be trialed in combination.

Most viruses adapt and mutate to become resistant to antiviral therapy and this can affect patient and disease management. This is exemplified by viruses including human immunodeficiency virus type 1, hepatitis C virus, and influenza A which have all shown the ability to develop resistance during single drug use therapies (*13–16*). Currently, there are no reports of circulating RDV-resistant strains of SARS-CoV-2. We are reliant on models based on studies in murine hepatitis virus (MHV), severe acute respiratory syndrome virus (SARS-CoV) and Ebola virus (EBOV) (*17–19*) in order to predict the amino acid residues that could, if mutated, confer drug resistance. Given the global threat presented by SARS-CoV-2, it is important to determine whether SARS-CoV-2 can become resistant to RDV, identify which mutations confer resistance, monitor the emergence of such variants in the population and adapt treatments in Covid-19 patients.

After determining optimal culture conditions (Fig.S1), SARS-CoV-2_Engl2_ was passaged serially in either 1μM or 2.5μM RDV-supplemented media for 13 passages (SARS-CoV-2_Engl2_ was isolated in February 2020; Fig. S2). Viruses serving as controls were passaged in parallel in either DMSO or media to monitor for cell culture adaptation. We passaged SARS-CoV-2_Engl2_ in parallel in 24 distinct cultures with different selective pressures (4 different conditions and 2 different virus inputs; Fig. S2). We monitored for cytopathic effect (CPE) during passaging of the cultures. CPE was observed in 7 of the 12 lineages passaged in RDV, with the loss of 5 lineages between p1 and p4 (Fig. 1A). There was general adaptation of the viruses to VeroE6 cells with an increase in overall viral titers by 0.5 to 1 log_10_ (Fig. 1A) as well as a change in plaque phenotype (Fig. S3A) after 13 passages. Next, the replication kinetics and change in RDV IC_50_ of a subset of passaged virus populations (Rem2.5p13.5, DMSOp13.5 and Mediap13.4) were assessed. Rem2.5p13.5 alone actively replicated in the presence of 7.5μM RDV (Fig. 1B). Although, titers in the presence of RDV were lower than those grown in the absence of RDV. Titers of control viruses, DMSOp13.5 and Mediap13.4 were consistently 5 log_10_ lower when cultured in the presence of RDV (Fig.1B). The Rem2.5p13.5, DMSOp13.5 and Mediap13.4 lineages displayed similar replication kinetics when cultured in the absence of RDV (Fig. 1B). When RDV sensitivity was assessed in VeroE6-ACE2 cells, Rem2.5p13.5 displayed a 2- to 2.5-fold increase in IC_50_ over a range of virus inputs in comparison with DMSOp13.5, and Mediap13.4 (Fig.S3B). The partial resistance to a nucleoside analogue was specific for RDV, as we observed a minimal change in IC_50_ of a second nucleoside analogue (EIDD2801), when comparing Rem2.5p13.5 (IC_50_ ~9.14μM) to SARS-CoV-2_Engl2_ (IC_50_ ~8.92μM) (Fig. 1C and Fig. S4).

**Fig. 1.**
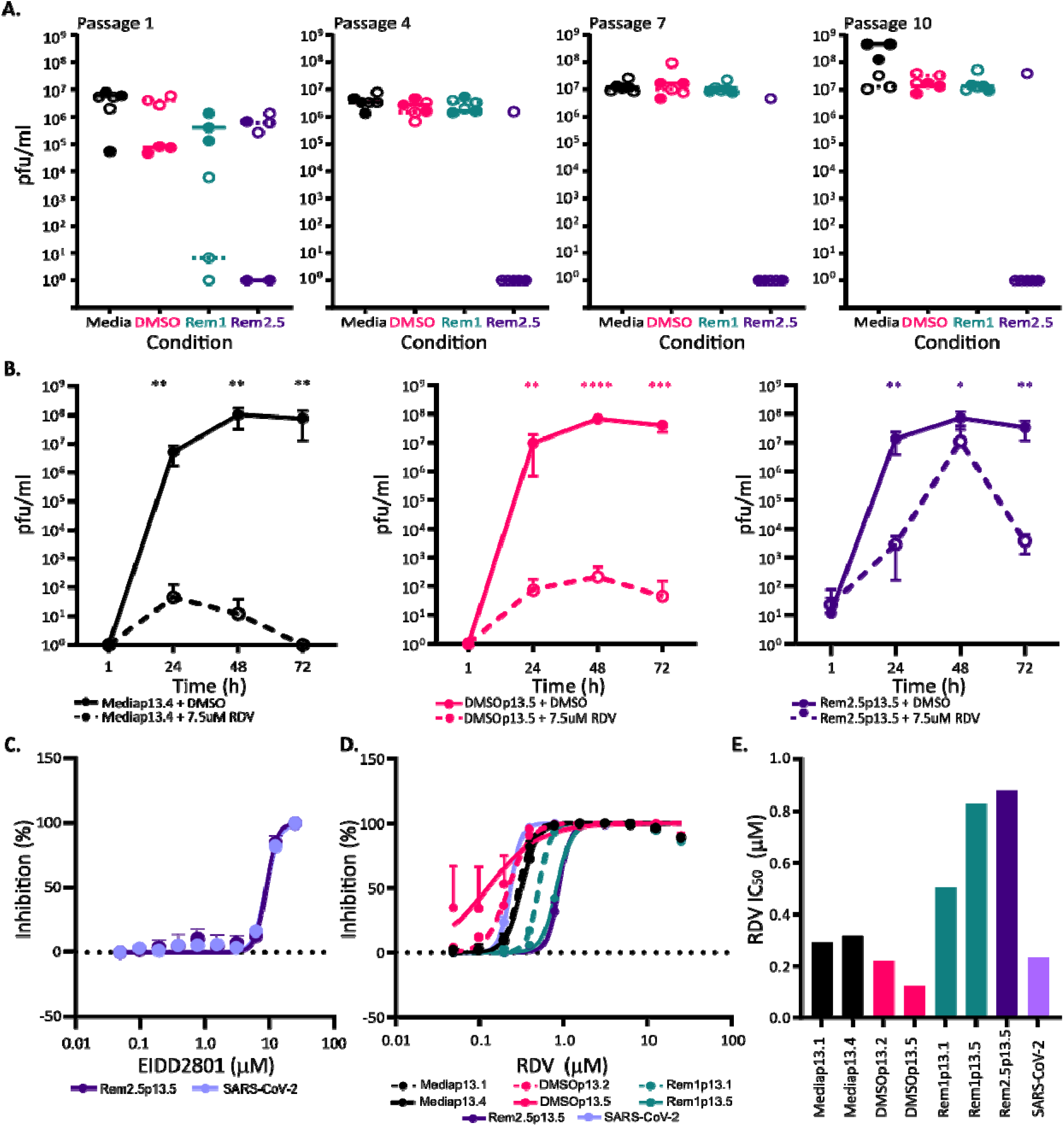
Continuous passage of SARS-CoV-2_Engl2_ in RDV selects for partial resistant populations. **A.** Virus titers (pful/ml) at p1, p4, p7 and p10. 6 lineages per condition and two different virus inputs; 1000 pfu (solid circle) and 2000 pfu (open circle). Median for each is shown. **B.** Virus growth kinetics in VeroE6 in the presence (dashed line) or absence (solid line) of 7.5μM RDV for 3 different virus populations. Data is from 2 independent experiments with 3 replicates. Error bars represent SEM. Unpaired t-tests (Holm-Šídák method; *,P< 0.05; **,P< 0.01; ***,P< 0.001. ****, P<0.0001). **C.** EIDD2801 dose dependency curve. EIDD2801 treated VeroE6-ACE2-TMPRSS2 infected with 8400 pfu/ml of each virus. **D.** RDV dose dependency curves determined in A549NPro-ACE2 infected with 8400 pfu/ml of each virus **E.** Bar graph of RDV IC50 for different viruses in A549NPro-ACE2 with 8400 pfu/well. For all panels, error bars represent SEM.

Subsequent analyses identified a second lineage, Rem1p13.5 with reduced sensitivity to RDV (Fig. 1D). The IC_50_ of Rem1p13.5 (~0.828μM) was comparable to Rem2.5p13.5 (~0.8281μM) and corresponded to a 3.5- to 3.7-fold increase from the parental virus (IC_50_~0.233μM). The RDV IC_50_ for virus passaged in either media alone (IC_50_~0.293-0.3159μM) or DMSO (IC_50_~0.124-0.221μM) corresponded with IC_50_ for the parental stock virus (Fig. 1D & 1E). The changes in RDV sensitivity paralleled those previously reported for MHV, SARS-CoV and EBOV resistant viruses (*3, 18*).

Direct comparison of the consensus sequences from all the passaged stocks with the original SARS-CoV-2_Engl2_ sequence revealed two fixed non-synonymous mutations in lineages with decreased RDV susceptibility in two independently generated populations (Rem1p13.5 & Rem2.5p13.5). These mutations were not present in either viruses passaged in absence of RDV, or the input virus (SARS-CoV-2_Engl2_) or SARS-CoV-2_Wu1_ (DataFileS1). The first mutation was identified as glutamine to aspartate at amino acid 802 (E802D) in the RNA-dependent RNA polymerase (RdRp) NSP12 (Fig. 2A). A glutamate at this position is highly conserved between all betacoronaviruses including SARS-CoV, MERS-CoV and unclassified sarbecoviruses (Fig.2B; DataFileS2). The E802 mutation occurs within the palm sub-domains (T680 to Q815; Fig.2A) and in proximity to amino acids predicted to interact with newly synthesized RNA (C813, S814 and Q815 (*20);* Fig.2A). We propose that the E802D mutation results in minor structural changes which reduce in steric hinderance in the region (Fig.2A), thereby influencing binding of nt+3 during synthesis of template RNA and allowing elongation when the active form of RDV is incorporated into the RNA. The mutation identified in NSP12 differs from amino acid residue involved with decrease RDV sensitivity in other betacornonaviruses, (MHV, SARS-CoV & MERS-CoV), and EBOV and predicted sites in SARS-CoV-2 (*17–19*).

**Fig. 2.**
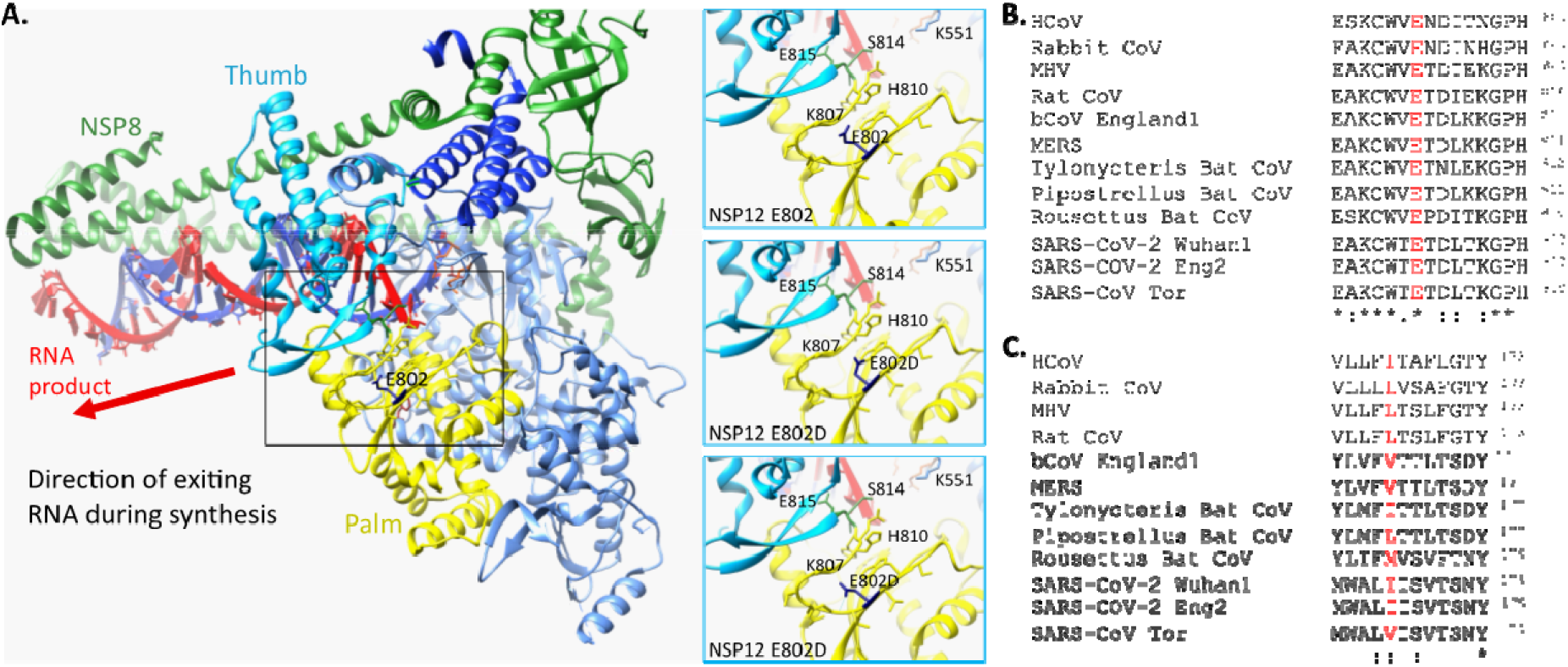
Common mutations in partial RDV resistance populations. **A.** Location of E802 within structure of SARS-CoV-2 NSP12 in association with NSP7 and NSP8 (PDB ID 6YYT). Three focused panels are WT (upper) and two potential confirmations of E802D. H-bonds are indicated by light blue line. **B.** NSP6 I168 amino acid is not conserved across coronaviruses. **C.** Conservation of E802 amino acid across coronaviruses. Accession numbers for the coronavirus sequences are in the materials and methods.

The second mutation was an isoleucine to threonine substitution (I168T) in NSP6, a highly conserved protein involved in restricting autophagosome expansion (*21*). This site is not highly conserved across coronaviruses with either an isoleucine (SARS-CoV-2) or valine (SARS-CoV & MERS-CoV) or leucine (MHV) in this position (Fig.2C; Data File S2). We predict that the mutation may alter the structure of the transmembrane and extracellular domains (Fig.S5A).

To ascertain whether a mutation of NSP12 E802 was sufficient to mediate partial RDV resistance, we introduced either an E802D or E802A mutation at this site into the backbone of SARS-CoV-2_Wu1_ and recovered infectious virus using a reverse genetics system. While unlikely to play a role, we also recovered virus with I168T mutation in NSP6 either alone or in combination with the NSP12 mutations (E802D or E802A). There were no significant differences observed in virus replication due to the mutations. All rescued virus mutants replicated similarly to the parental rSARS-CoV-2 in human lung cells, Calu-3, with similar replication kinetics and achieving similar peak virus titers (Fig. 3A). Both the E802D and E802A mutations in NSP12 recapitulated partial resistance observed in the virus populations continually passaged in RDV (Fig. 3B). We observed a 2.47-to 2.097-fold change in RDV IC_50_; from 2.298μM for rSARS-CoV-2 to 5.676μM and 4.818μM for the E802D and E802A mutants, respectively (Fig. 3B; Table. S1). This change in RDV sensitivity was evident over a range of virus inputs for both NSP12 mutants (Fig. S6A). NSP6 I168T substitution did not confer decreased sensitivity to RDV (Fig. 3B), with the IC_50_ calculated comparable to rSARS-CoV-2 (Table S1).

**Fig. 3.**
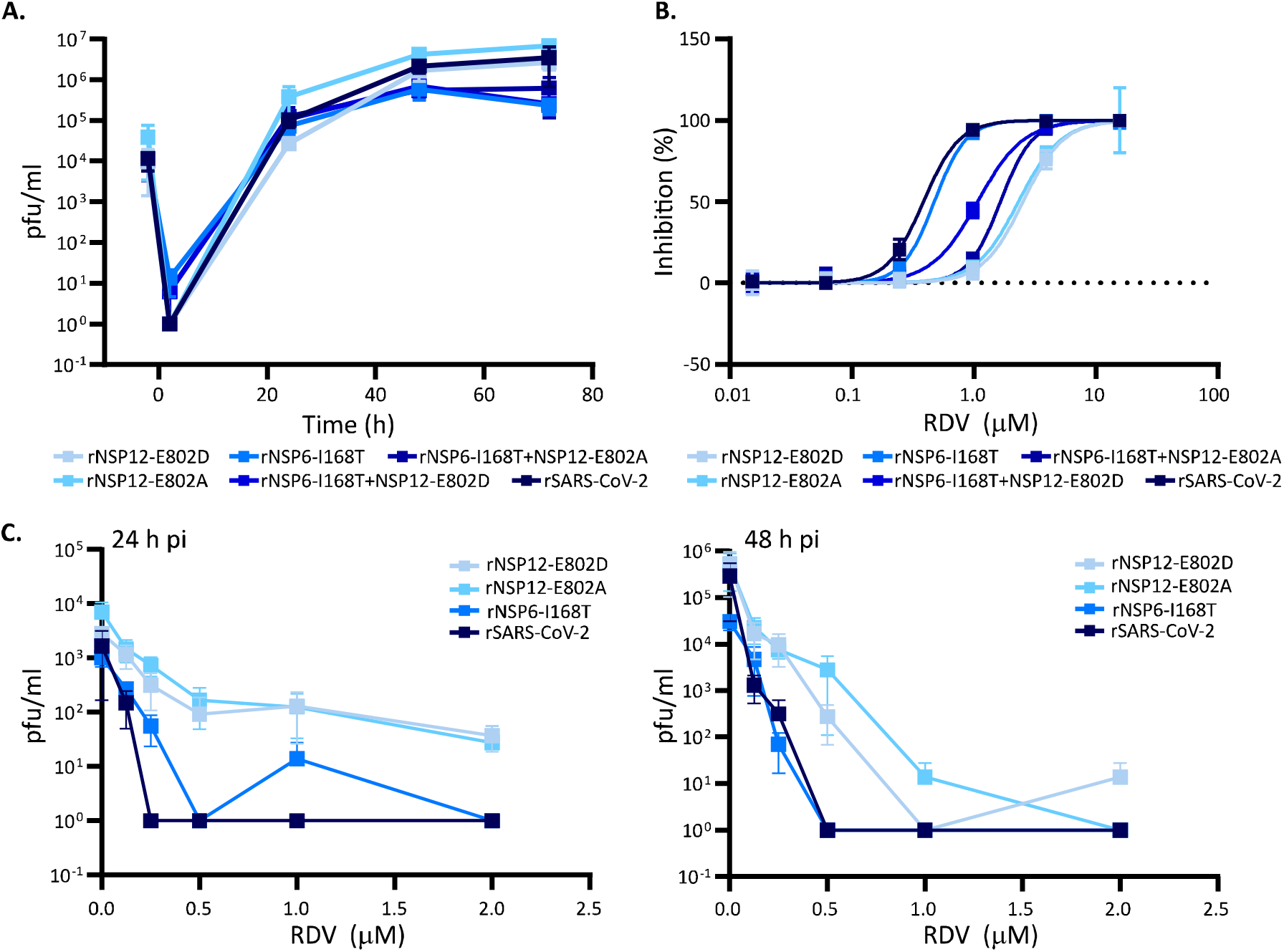
NSP12 E802 mutation recapitulates change in RDV susceptibility. All viruses were derived by reverse genetics and have a SARS-CoV-2_Wu1_ backbone with specific point mutations as indicated. All error bars are SEM.**A.** Virus replication kinetics of rescued viruses with single mutation in either NSP12, NSP6 or both in Calu-3. Data is from 3 independent experiments with 3 replicates, there was no significant difference between growth of the mutants versus the wild-type rSARS-CoV-2. **B.** RDV dosedependent inhibition for each mutant virus. Mutations in NSP12 decrease the sensitivity to RDV. **C.** RDV dose effect on virus titers at 24 h (left) and 48 h (right). Data from 2 independent virus stocks with 2 replicates except for rSARS-CoV-2 and rNSP12-E802A

Indeed, viruses bearing both the NSP12 and NSP6 mutations were more sensitive to RDV in comparison to NSP12 single mutant viruses (rNSP6-I168T+NSP12-E802D, IC_50_ 3.728μM; rNSP6-I168T+NSP12-E802A, IC50 3.096μM). Importantly, introduction of NSP6 and/or NSP12 mutations did not significantly affect sensitivity to EIDD2801 (Table. S2). These data confirm results obtained with other viruses indicating EIDD2801 sensitivity was not influenced by mutations conferring decreased RDV sensitivity (*18, 22*). We further assessed the anti-viral activity of RDV in Calu-3. While a dose-dependent reduction in titer for all viruses was observed, rNSP12-E802D and rNSP12-E802A titers were consistently higher than wild-type and rNSP6-I168T at 24 and 48h (Fig. 3C). Interestingly, at 24h pi, a slight shift in an increase rNSP6-I168T infectious titer was observed in comparison with wild type, though this effect disappeared by 48h.

We next examined the available SARS-CoV-2 genome sequences in the CoV-GLUE database (n=242865 as of January 2021) and searched for sequences with replacements at NSP12 E802 and NSP6 I168. Only 8 viral sequences in total were identified with a mutation at E802; four sequence had E802A (3 sequences sampled in May 2020 from the same geographic region) while four sequences with E802D were geographically dispersed. As one of these sequences, hCoV-19/Scotland/CVR2716/2020 was isolated from a patient who was not treated with RDV, these suggests mutation of E802 can be selected in the community in the absence of drug selection. The observed global frequency of the E802 substitutions was the same as mutations at either NSP12 F480 or V557; sites known to confer partial RDV resistance in other coronaviruses (*18*). There were a handful of sequences with changes at either F480 (n=5) or V557 (n=6). Replacement of NSP6 I168 occurred in 33 sequences with isoleucine replaced with threonine, valine, leucine or methionine. These data indicate that in absence of selective pressure, mutations of either NSP12 E802 or NSP6 I168 are rare events. However, the identification of these sequences in the genome databases demonstrate that these viruses are viable and could potentially acquire a resistant phenotype when a selective pressure is applied.

To our knowledge there are no reports identifying signatures within the genome of SARS-CoV-2 which lead to resistance (or partial resistance) to RDV. We should consider that our partially-resistant RDV populations arose rapidly with a fixed lineage within 4 passages rather than 23 to 30 passages as observed for MHV and Ebola, respectively (*18, 19*). The potential for resistance to occur in RDV non-responding patients may be a an issue that needs to be examined in order to discern whether it is due to a genomic mutation or drug tolerance by synchronization (*23*). The change in sensitivity to RDV was similar to single NSP12 mutation in either MHV or SARS-CoV (*18*) but lower than EBOV (*19*).

We observed our SARS-CoV_Engl2_ RDV-resistant viruses and the reverse-genetic derived SARS-CoV-2_Wu1_ NSP12 mutants increased the IC_50_ by at least 2-fold regardless of the cell type used for the experiments. Thus, we are confident that the change in IC50 was not due to cellular drug metabolism or differences in virus entry and replication between wild-type and RDV-resistant viruses. We also noted that the cell-culture adaptation in viruses passaged in the absence of RDV resulted in a shift in IC_50_ in VeroE6 based assays in comparison with input SARS-CoV-2_Engl2_ (Fig.S4) but this shift was not as predominant as the RDV-selected viruses. We hypothesize that this was due to more efficient virus entry and spread as many of the mutations observed occurred within the spike protein (see below). Difference in IC50 due to adaptation, availability of receptors and ability to metabolize RDV is widely acknowledged (*3, 24*).

We next focused on those mutations arising in the *in vitro* passaged virus populations that were likely not directly linked to RDV resistance. The consensus sequences of all the passaged stocks displayed a total of 41 distinct non-synonymous mutations and 10 synonymous mutations across the genome compared to the parental SARS-CoV-2_Engl2_ sequence (Fig. 4). Importantly, we did not observe any previously identified mutations in the proof-reading ExoN (NSP14) that would change the sensitivity of the virus to RDV (Fig. 4). Deletions of ExoN have been demonstrated to increase RDV sensitivity for other coronaviruses (*18*). While there was clear positive selection pressure across the entire genome (Table S3), there were no major differences in the number of mutations that accumulated in any specific population, and in the ratio or type of transition vs transversion change (Fig 4B & S5B). Although, Rem2.5p13.5 displayed a slight elevation in non-synonymous changes (Fig 4C), we are unable to draw conclusions on the effect of RDV concentration on virus mutation rate due to recovery of an insufficient number of populations selected in RDV.

**Fig. 4.**
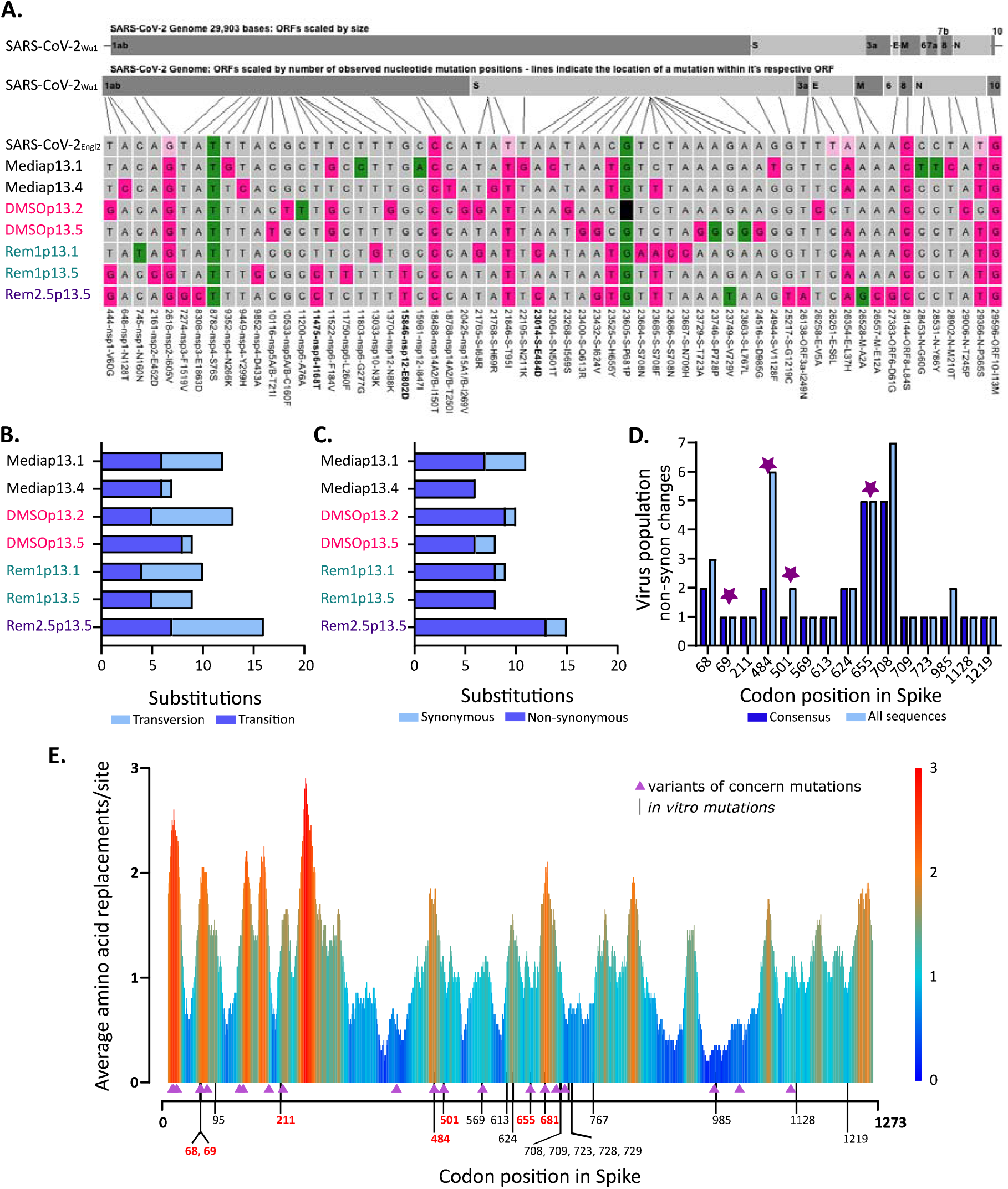
Sequence analysis of partial resistance RDV populations. **A.** Alignment of serially passaged viruses and SARS-CoV-2Engl2 to SARS-CoV-2Wu1. Non-synonymous (pink) and synonymous (green) changes from Wuhan-1 are highlighted. Light pink are sites fixed at 50% in SARS-CoV-2_Eng2_ and black box is a deletion mutation in serially passaged virus. Positions of mutations are indicated, and mutations only found in RDV selected populations are in bold. **B**. Synonymous vs non-synonymous changes observed in continually passaged virus populations compare to input SARS-CoV-2_Eng2_. **C.** Transversion vs transitional changes observed in continually passaged virus populations compare to input SARS-CoV-2Eng2. **D.** Number of *in vitro* passaged viruses with non-synonymous changes in Spike in comparison to SARS-CoV-2Engl2. Mutation fixed in the consensus genomes (dark blue) are compared to the total number of viruses with evidence of the mutation at sub-consensus levels (light blue). Amino acid residues in common with the emerging variants of concern (UK B.1.1.7, Brazil P.1; and South Africa B.1.351) are highlighted by a star. **E.** Worldwide diversity of Spike protein sites of circulating SARS-CoV-2 variants. The average number of different substitutions at each codon is calculated along a 20 amino acid residues wide sliding windows. Data was calculated using only substitutions observed in a minimum of 5 sequences from the publicly available SARS-CoV-2 genomes (n=1384). The position of the amino acid substitutions in the *in vitro* passaged viruses are indicated at the bottom, residues are red are shared with variants of concerns, black are specific to *in vitro* virus, residues in italics were synonymous. Mutations in amino acid residues that are also mutated in the variants of concern (UK B.1.1.7, Brazil P.1, & South Africa B.1.351) are shown in purple triangles.

Most of the mutations (22 mutations) occurred within the spike (S) open reading frame. Unlike other studies (*25–27*), the furin-like cleavage site was preserved in all but one of the passaged populations; DMSOp13.2 displayed a 24nt deletion of the entire furin-like cleavage site at a high but unfixed frequency of 77%. Further comparative analysis with the SARS-CoV-2_Wu1_ sequence identified a further 2 synonymous and 8 non-synonymous mutations present in the original SARS-CoV-2_Engl_2 population (Fig. 4A). SARS-CoV-2_Engl2_ was a 50:50 mix of two virus populations with 5 of the mutations present at a frequency ~50%, all but one of these became fixed in all passaged populations by p13 (Fig. 4A).

Importantly, in our *in vitro* passaged viruses we observe substitutions at the same sites within Spike (H69, E484, N501, H655, P681) that were also identified in the emerging SARS-CoV-2 variants of concern (B.1.1.7: Δ69/70, N501, P681; P.1: E484, N501Y, H655Y; B.1.351: E484K, N501Y) (Fig.4). Except for synonymous P681P, these substitutions were not present in SARS-CoV-2_Engl2_ (Data File S1). Of note, while the E484 mutation appeared in the consensus sequence of Rem2.5p13.5 and Remp1p13.1 (Fig.4A & 4D), it was present at a frequency of 20-40% in all the other viruses with the exception of DMSOp13.2 (Fig. 4D). The N501 substitution was present in one virus at consensus (Mediap13.1) and also present in the subconsensus of a second (Rem2.5p13.5) (Fig. 4D). It is important to stress these emerging variants of concern, collectively, share a combination of three amino acid mutations in Spike: E484, N501 and K417, with N501 common to all. Two of these mutations are observed in our *in vitro* evolution studies. The probability of large overlap (5 codons) between the substitutions observed *in vitro*, and variants of concern defining mutations without a common selective pressure driving convergence, was exceptionally small (P=3.1×10^−5^; Fig. S8). This demonstrates commonality in the fitness landscape that these *in vitro* populations and the circulating lineages are evolving under. We further examine the global distribution of all circulating amino acid replacements within Spike to determine whether our *in vitro* substitutions occurred within hot spots for change. There were 1384 replacements observed in a minimum of 5 sequences (n=242865 sequence up dated 14^th^ December 2020), many of these were clustered into certain regions within Spike, creating visible hot spots of diversity (Fig. 4E). For example, the window surrounding amino acid E484 appears to be a relative hot spot for replacement. These observations underline the plasticity of the SARS-CoV-2 genome and suggests independent emergence of geographically different variants sharing common mutations have not necessarily occurred due to immune-based selection pressure. Our data shows these mutations arise *in vitro* in the absence of any immune selection.

In summary, we have identified in *in vitro* evolution studies a genome signature in SARS-CoV-2 which allow replicative advantage in the presence of RDV. In the US, RDV treatment is currently prescribed to at least half of all hospitalized SARS-CoV-2 patients (*28*). Our data demonstrates that selection of RDV resistance in SARS-CoV-2 can occur but there is no evidence of global spread of RDV-resistant strains. In addition, we have shown that key amino acid residues that have been identified in emerging variants of concerns in three different continents can occur *in vitro* in the absence of immune pressure. Overall, our study offers new perspectives for the surveillance of new SARS-CoV-2 variants and the clinical management of patients treated with RDV.

## Supporting information

Supplementary DataFile S1

Supplementary DataFiles S2

## Acknowledgments

We would like to thank all the global researchers who shared their data on GISAID (https://www.gisaid.org),COVID-19 Genomics UK Consortium (COG-UK) and Kyriaki Nomikou, Jenna Nichols, Yasmin Parr, Lily Tong and Natasha Johnson from the CVR Genomics Team.

## Author contributions

MES, AMS, ECT & AK conceived the project; MES, AMS, AM, SJW, AHP & AK designed the methodology; MES, AMS, AM, SW, MT, GL, RMP, & AW undertook the experiments; bioinformatic analysis was performed by RJO, OML & MES;ASF, DM, RJO & ECT were responsible for NGS and data was curated by MES, AMS, OML & RJO. RJO, OML & MES Writing-original draft preparation by MES, AMS, RJO, BJW, MP & AK and visualization by MES, AMS, OML & RO. All authors reviewed the final manuscript. Project administration: MES, AMS, AK and funding was acquired by MP, BJW, AK, SJW, AHP, ASF.

## Data and materials availability

SARS-CoV-2Engl2 was supplied under a MTA between The University of Glasgow and Public Health England. Consensus sequences and raw FASTQ files have been uploaded to GenBank under BioProject number PRJNA692078 and will be released upon publication. We used the publicly available CoV-Glue database (http://cov-glue.cvr.gla.ac.uk/#/home) to examine for replacements at specific sites observed in the GISAID hCoV-19 sequences. All data is referred to in the main text or the supplementary materials is available.

## Supplementary Materials

Materials and Methods

Figures S1-S7

Tables S1-S3

Data Files S1-S2

References (*##-##*)

