## Supplementary material for "*In vitro* evolution of Remdesivir resistance reveals genome plasticity of SARS-CoV-2": M&M, Supp Text, Figures and Tables

**This PDF file includes:**

Materials and Methods

Supplementary Text

Figs. S1 to S8

Tables S1 to S3

Captions for Data S1 to S2

References

Materials and Methods

**Cell culture.** African Green monkey kidney cells VeroE6 expressing ACE-2 (VeroE6-ACE2) alone and with TMPRSS2 (VeroE6-ACE2-TMPRSS2), BHK-21, and A549NPro expressing ACE-2 (A549NPro-ACE2) were grown in DMEM-Glutamax supplemented with 10% fetal calf serum (FCS; Gibco) and non-essential amino acids (NEAA; Gibco). VeroE6 were a kind gift from Professor M. Bouloy (Institut Pasteur, France), BHK were sourced from the ATCC (ATCC CCL-10) and, VeroE6-ACE2, and VeroE6-ACE2-TMPRSS2 have been described elsewhere (*29*) A549NPro cells (kind gift from Prof. Rick Randall) were transduced to stably express ACE-2 (A549NPro-ACE2)as described (*29*) Calu-3, a human lung epithelial cell line was grown in DMEM Glutamax, supplemented with 20%(v/v) FCS and NEAA was sourced from the ATCC ([ATCC HTB-55)](https://www.lgcstandards-atcc.org/products/all/HTB-55.aspx). All cells were maintained at 37 °C in 5% (v/v) CO_2_, humidified incubator.

**Viruses.** All initial work with SARS-CoV-2 was undertaken with the clinical isolate SARS-CoV-2_Engl2_ strain (hCoV-19/England/02/2020 (England-02), GISAID accession: EPI_ISL_407073) kindly supplied by Public Health England. Validation of the mutations was undertaken in rescued SARS-CoV-2 based on Wuhan-Hu-1 (MN908947) sequence. All virus stocks were grown in VeroE6 cells as described, supplemented with 4%(v/v) FCS, and for the drug screens regardless of cell type the final concentration of FCS was 2%(v/v). For virus growth curves, FCS concentration was decreased to 2%(v/v) in Vero E6 while in Calu-3 FCS concentration was 10%(v/v).

**Plaque assays.** SARS-CoV-2 titers were determined by plaque assay in either VeroE6 or VeroE6-ACE2-TMPRSS2. Cells were seeded at 2.5 x 10^5^ cells/well in 12-well plates and plaque assays as described (*29*).The plates were fixed in 8% formaldehyde in PBS, stained with Coomassie Blue staining solution (0.1% [w/v] Coomassie Brilliant Blue R-250; 45% [v/v] methanol; 10% [v/v] glacial acetic acid) and imaged with a photo scanner (Epson Expression 1680 Pro).

**Chemical inhibitors.** RDV (GS-5734; HY-104077) and EIDD-2801 (HY-135853) were purchased from MedChemExpress, diluted to final concentration of 10 mM stock solutions in 100% (v/v) dimethyl sulfoxide (DMSO, Sigma) and stored at -80C~~.~~

**Plasmids, virus rescue and validation of RDV resistant viruses.** Reverse genetics plasmids used to rescue SARS-CoV-2_Wu1_ have been described previously (*29*). Briefly, specific point mutations in SARS-CoV-2 cDNAs in fragment 3 (nt11475 T to C) and fragment 5 (nt15846 either A to T or A to C) were synthesized by Biobasic and assembled into the full-length molecular clone to give pCCI-4K-SARS-CoV-2-NSP12_E802D, pCCI-4K-SARS-CoV-2-NSP12_E802A, pCCI-4K-SARS-CoV-2-NSP6_I168T, pCCI-4K-SARS-CoV-2-NSP6_I168T-NSP12_E802D-NSP6, pCCI-4K-SARS-CoV-2-NSP6_I168T-NSP12_E802D-NSP6. The pCCI-4K based plasmids containing the full-length SARS-CoV-2_Wu1_ sequence with the mutations were propagated in TransforMax™ EPI300™ Electrocompetent *E. coli* and purified by using a Macherey Nagel Endotoxin free plasmid purification kit. For virus rescue, BHK-21 cells were transfected with 3 ug of plasmids using Lipofectamine LTX reagent as per manufacturer’s recommendation. Cell culture supernatant was harvested 48 and 72h post transfection (p0) and used to infected VeroE6 to generated p1 stock of the recovered viruses.

**SARS-CoV-2 compound sensitivity studies and IC_50_ calculations.** To test for a change in sensitivity to RDV of our continually passaged SARS-CoV-2_Eng2_ viruses and reverse genetic derived viruses (rNSP12-E802D, rNSP12-E802A, rNSP6-I168T, rNSP6-I168T-NSP12-E802D, rNSP6-I168T-NSP12-E802A and rSARS-CoV), two different compound screen layouts were tested (Fig. S7) Cells were seeded at 1.2 x 10^4^ cells/well for VeroE6, VeroE6-ACE2, and VeroE6-ACE2-TMPRSS2 and 1.3 x 10^4^ cells/well for A549NPro-ACE2 cell lines in a 96-well format. Two-fold serial dilutions of RDV starting at either 25 or 50 μM were added to the cells and incubated for a minimum of 2h prior to addition of virus. Depending on plate layout, either a set amount of virus or 2-fold serial dilutions of the virus were added, and plates incubated in a 37°C humidified incubator in 5% CO_2_ for 72h. For the mixed array plate layout, each drug dilution and virus combination were in triplicate on each plate and repeated twice. Plates were fixed in 8% (w/v) formaldehyde for 1h and stained in Coomassie Brilliant Blue solution. Washed and airdried plates were scanned using the Celigo (Nexcelcom) and the total intensity of each well was determined. The total intensity values from the Celigo scan were used to calculate the IC_50_ values using Prism GraphPad v8 and v9. Analysis programs used included transformation of concentration (log_10_), normalization of transformed data and non-linear regression (log(inhibitor) vs. normalized response -- variable slope).

**Selection of RDV resistant viruses.** Continual passages of SARS-CoV-2 were performed in the presence of RDV in VeroE6 cells. Assays were undertaken in a 12 well plate with each condition in triplicate; two RDV concentrations (1μM and 2.5μM) and two virus inputs (1000 & 2000 pfu) were used, and controls for cell culture adaption of the virus including media alone or 0.2% (v/v) DMSO were undertaken in parallel (Fig.S3). These concentrations were kept constant throughout the experiment. Time between each amplification was dependent of visible CPE from passage 1 to passage 5 (p1 to p5), there was up to 7 days between each amplification. For these passages, 100 μl of each amplification was used to generate the subsequent passage. It became evident after passage 4 (p4) that some of the conditions used resulted in loss of infectious virus. After passage 5, 10 μl of culture media was used for passaging the virus in wells that showed evident CPE. Where CPE was not evident, 100μl of cell culture was blindly passaged. After each passage the cell media was harvested and stored at -80°C. Samples from p4, p7, p10 and p12 were also harvested for titration and cell monolayer was harvested in TrizolLS for future analysis. Prior to sequencing, each condition was tested to determine its ability to grow in the presence of RDV.

**Virus replication kinetics.** To compare the replication kinetics of the parental stock, cell culture adapted and RDV resistant mutant SARS-CoV-2, monolayers of VeroE6 cells were treated with 7.5μM RDV and infected at a MOI~0.1. The inoculum was removed after 1h, cells washed and fresh media containing either DMSO or RDV added. Growth curves in VeroE6 were undertaken twice in triplicate. rSARS-CoV-2_Wu1_ and mutant viruses virus growth was also assessed in Calu3 cells. For this, Calu-3 were seeded 7 days prior to infection, media was changed every 2-3 days prior to infection. The cells were infected with a MOI~0.01 (based on VeroE6-ACE2-TMPRSS titres) for 1h, inoculum then aspirated, cells washed, and fresh media added. All infections for virus growth curves were asynchronous. Cell culture media from each well were collected at 24, 48 and 72h post infection (p.i.) and cell-free virus titers were determined by plaque assay in VeroE6 cells. Each experiment was performed at least thrice with a triplicate of the virus each time (mutant viruses n = 3, wt n=6). Virus titers at each time were determined by plaque assay as described above.

To assess the influence of RDV on the growth of rSARS-CoV-2 and mutant viruses, Calu-3 were seeded 7 days prior to infection in a 24 well format (2 x 10^5^ cells/well), media was changed every 2-3 days. Two hours prior to infection 2-fold serial dilutions of RDV from 2 μM to 0.125μM. Cells were infected at a MOI of 0.01 asynchronously for 1 h, inoculum aspirated, and media containing the appropriate amount if RDV added. Samples were taken 24 and 48h p.i. Titers were determined by plaque assay. Data from 2 independent virus stocks with 2 replicates except we wre missing one of the replicates for wild-type and a 48h replicated for rNSP12-E802A at 48 h.

**RNA purification from virus stock.** Clarified supernatant from p13 was added to TrizolLS and total RNA was extracted as described (*30*) for sequence analysis.

**Sequence of virus stock using Illumina MiSeq.** Sequencing libraries composed of overlapping amplicons across the SARS-CoV-2 genome were prepared utilizing the ARTIC Network protocol (<https://artic.network/ncov-2019>). Amplicons were paired-end sequenced (2x250 nt) on an Illumina MiSeq as described previously (*31*). Reads were trimmed with trim_galore (http://www.bioinformatics.babraham.ac.uk/projects/trim_galore/) and then mapped with Burrows–Wheeler Aligner (*32*) to the SARS-CoV-2_Wu1_ (MN908947) genome, followed by amplicon primer trimming and consensus calling with iVar (*33*) using a minimum read coverage of 10. Consensus sequences and raw FASTQ files have been uploaded to GenBank under BioProject number PRJNA692078.

**Modelling of mutations in SARS-CoV-2 nsp12.** UCSF Chimera (*34*) was generate the ribbon structure and rotamer function (*35*) used to model the mutations into SARS-CoV-2 _Wu1_ NSP12 (PDB 6YYT(*36*)). Protter (*37*) was used visualize the effect of the I168T mutation on organization of the transmembrane domains in NSP6.

**Statistics and bioinformatic.** GraphPad Prism v8 & v9 software (La Jolla, CA, USA) was used to undertake statistical analysis (transformation, normalization, non-linear regression) and generate figures. EMBL-EBI Clustal omega (*38*) was used to align the sequence of NSP6 and NSP12 from SARS-CoV-2 Wuhan1 (MN908947), betacoronavirus England1 (bCoV England1; YP009944302 & YP009944297.1), SARS-CoV-2_Eng2_, SARS-CoV Tor (NP_828849.7 & YP009944371.1), human coronavirus (hCoV, HKU, 1YP459941.1 & YP009944274.1), middle eastern respiratory virus (MERS; YP009047223.1 &009047234.1), murine hepatitis virus (MHV; YP009924352.1 & 009915678.1), rat coronavirus (Parker, YP009924378.1 & YP_009924378.1), Rousettus bat coronavirus (HKU9; YP009924393.1 & YP009924388.1), rabbit CoV (HKU14; YP009924419.1 & YP009924414.1), Tylonycteris bat CoV (HKU4 YP009944320.1 & YP009944315.1), and Pipostrellus bat CoV(HKU5; YP009944349.2 & YP009944344.1). The online database CoV-GLUE (<http://cov-glue.cvr.gla.ac.uk/>(*39*)) was used to monitor for replacement in the SARS-CoV-2 genome. The database was accessed on 10th of January 2021 (n = 242865). The dN/dS ratios were calculated for each sample using all ORFs concatenated together, as well as the Spike and entire ORF1ab sequences individually (with respect to the SARS-CoV-2Eng2 sample to monitor within passage evolution), using the program SNAP (<https://www.hiv.lanl.gov/content/sequence/SNAP/SNAP.html>) (*40*)

The code used to calculate the null distribution and the probability as well as the global distribution of the mutations within in Spike can be assessed at [https://github.com/omaclean/CV19IV/](https://github.com/omaclean/CV19IV/blob/main/permutation.R).

The receptor binding domain of Spike is from amino acids 319-541: B1.351 and P1 had 3 mutations in the RBD (K417N/T, E484K and N501Y) while B.1.1.7 only had one (N501Y). We used the following amino acids replacements and deletions in Spike for each lineage based COG-UK defined changes for each lineage; B1.351 (L18F, D80A, D215G, R246L, K417N, E484K, N501Y, A701V), P.1 (L18F, T20N, P26S, D138Y, R190S, K417T, E484K, N501Y H655Y, T1027I), B.1.1.7 (Δ69-70, Δ144, N501Y, A570D, P681H, T716I, S982A, D118H). We further included the substitution observed in P1 (V1176F) and deletion in B1.351 (Δ242-245). There is phylogenetic uncertainty if V1176F was present in the ancestor preceding the formation of the P.1 lineage, the deletion (Δ242-245; like other defining mutations) in B.1.351 is polymorphic. We erred on the side of inclusivity in the face of this uncertainty, making our analysis more conservative. All sequences available sequences for SARS-CoV-2 were assessed from CoV-Glu (n= 242865, last update 14^th^ December 2020).

Supplementary Text

**Adaptation of SARS-CoV-2 to VeroE6**

General adaption of the virus led to an over shift in the IC_50_ of the continually passaged virus in comparison to the input virus SARS-COV-2_Engl2_. This was clear with both the RDV IC_50_ and EIDD2801 IC_50_ for DMSOp13.5, and Mediap13.4 virus lineages (Fig. S4) determined in cell lines with a VeroE6 backbone. This was a 2.5 to 2.7-fold and 2.1-fold increase in RDV and EIDD2801 IC_50_ required to protect the cells against a dose of SARS-CoV-2_Engl2_ (Fig. S4) that would cause complete clearance of a 96-well. The RDV IC_50_ fold-change for the REM2.5p13.5 in these cells was 4.54 when compared to SARS-CoV-2_Engl2_ and was 1.64 to 1.78-fold higher when compared to the continually passaged viruses. This can be directly compared to the fold change of the RDV IC50 for the same viruses in A549NPro-ACE2 cells. The fold-change of viruses passaged in the absence of RDV was equivalent to SARS-CoV-2_Engl2._ Both DMSO adapted viruses were more sensitive to RDV while the media adapted were similar to SARS-CoV-2_Engl2_. There are other caveats that should be consider including metabolism of the antivirals to their active forms between the two cell types (*3*, *24*)

**Does RDVS cause SARS-CoV-2 to mutate?**

It is not an uncommon phenomena that nucleoside analogue cause higher mutation rates in RNA viruses (*41*, *42*). While there is no difference in the number of mutations that accumulate between the virus passaged in low concentration of RDV treated samples (Rem1p13.1 and Rem1p13.5) and the virus population grown in the absence RDV (Fig 4, DataFile S1). There is, however, a very clear difference in number of mutations accumulated Rem2.5p13.5 in comparison to all other passaged viruses. This may infer that higher concentration of RDV does increase mutation rate. We would suggest caution is used when making this conclusion as sample size for all groups was small and we have no replicates for viruses adapted in a higher concentration of RDV.

**Are the samples under selective pressure to evolve?**

Calculation of the dN/dS ratios of the concatenated genomes showed clear evidences of positive selective pressure in a majority of the continually passaged viruses regardless of the presence of RDV (Table S3). This is further supported when examining Spike and ORF1ab individually. As mentioned, this needs to be caveated with our small sample size, the high dN/dS ratios are influenced by the lack of observed syn mutations. (i.e. if 0 syn mutations and >0 non-syn mutation, then dN/dS will be infinity).

**Are the sites in Spike under adaptive pressure to change?**

We were surprised to observe the mutation of in our passaged viruses that were occurring in same location as those identified in the newly emerging variants from the UK, South Africa and Brazil that are associated with increased transmission (*43*). The current hypothesis suggests that these mutations are due to immune selection and/or pressure. Our data would suggest that SARS-CoV-2 can and does use positions to allow adaptation to very different conditions. These Spike mutations arose in our passaged viruses; I68R, H69R, T95I, N211K, E484D, N501T, I569S, Q613R, I624V, H655Y, P681P, S708N/F/F, N709H, T723A, P728P, V729V, D985G, V1128F, and G1219C. Not all mutations occurred in every lineage, and the frequency also varied between lineage. We are confident selective pressure occurring with substitution of E484 (10% to 80% fixed) and H655 (46 to 99% fixed in 5 populations) arising within all 7 passaged. While N501 mutation was in 2 lineages (16 to 61%). There are also synonymous changes occurring at both P681 and P728 rather than non-synonymous change associated with the circulating variants. Furthermore, we have a substitution at H69 rather than a deletion. The T95I and P681 changes were present in the input virus but otherwise the others have arisen through passage. A publication by Vega et al (*44*) highlighted that the *in vitro* mutation rate of SARS-CoV-2 in VeroE6 was negligible indicating that it was well-adapted for growth in cell culture. This would indicate that there is further selective pressure on these sites in Spike to change in independently evolving virus populations to provide an advantage.

To further examine the likelihood of the observed overrepresentation of the 21 variant of concern defining spike mutations in our *in vitro* passaged viruses, we simulated mutational events under a null distribution. This null distribution assumed that all codons were equally likely to be mutated within Spike (except for the start and stop codons; amino acid positions 1 and 1273 respectively). Each run involved 20 codons in Spike being sampled (without replacement- i.e. each codon could be mutated maximally once), and the number of sampled codons which overlapped the VOC sites were recorded. The distribution (plotted below) over 100 million simulations only produced five overlapping mutations 2981 times, giving a probability of 3.1x10^-5^ observing this level of overlapping evolution if the *in vitro* evolution was random. It is important to note that the null distribution was not impacted by the single deletions of amino acids 68 and 69 in the UK B.1.1.7 variant as mutations of these sites in the *in vitro* sequences required two mutations in the *in vitro* data.

We further investigated the global distribution of the mutations within in Spike to examine whether our *in vitro* substitutions occurred within hot spots of amino acid replacements in the global SARS-CoV-2 sequence database. In Figure 4E, each bar on the y axis represents a sliding window of 20 amino acids, counting the average number of unique amino acid replacements observed for each site in that window. The method we used does not incorporate information from the phylogeny and only counted unique observed states, as recorded in CoV-GLUE, using GISAID data up to the 14^th^ December 2020. For example, the N501Y replacement has occurred numerous times in parallel (<http://cov-glue.cvr.gla.ac.uk/#/project/replacement/S:N:501:Y>), but it will only be counted once in our analysis, though the N501T and N501S substitutions will be counted as additional replacement (<http://cov-glue.cvr.gla.ac.uk/#/project/replacement/S:N:501:T> ; <http://cov-glue.cvr.gla.ac.uk/#/project/replacement/S:N:501:S>) In order to avoid counting sequencing errors or strongly deleterious variants, amino acid replacements seen in fewer than five independent sequences on the CoV-Glu database were ignored. (For tree aware selection analyses, which incorporate observations of parallel mutation events, see <https://observablehq.com/@spond/revised-sars-cov-2-analytics-page>). Figure 4D shows clear overlap and shared clusters of substitution shared between variants of concern and the *in vitro* passaged virus within Spike.

It should be noted that reference stock of SARS-CoV-2_Engl2_ provided from PHE underwent two rounds of amplification in VeroE6 prior to use in our continual passage study (Fig.S2). Deep sequence of the amplified stock revealed diversification of the amplified virus away from the reference sequence. There were five derived non-reference mutations present at a frequency of ~50%, four of these five became fixed across all passaged populations by p13. (DataFileS1). In our analysis of substitutions of Spike, we have included T95I as a tissue culture derived mutation as it was at ~46% frequency in the input virus and fixed at 99 to 100 in all bar one virus (Fig. 4A).

Supplementary Figures

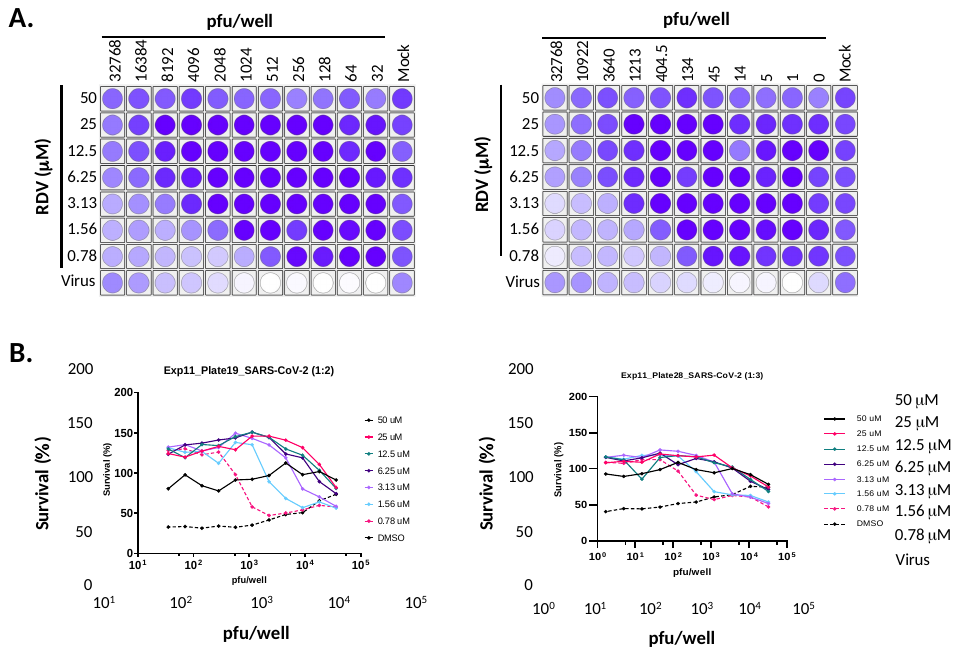

**Fig. S1. RDV mixed array inhibitor screen with SARS-CoV-2_Engl2_ in VeroE6 cells.** Two-fold serial dilutions of RDV from 50μM and either a two-fold (left) or three-fold (right) dilution of SARS-CoV-2. **A.** Heatmap of total intensity of the stained plated generated by the Celigo. Lightest colors indicate clearance of the monolayer. **B.** RDV dose dependency over a range of virus inputs. For each RDV concentration the survival (%) of the monolayer with different virus inputs per well is plotted. Values are from 1 replicate per condition.

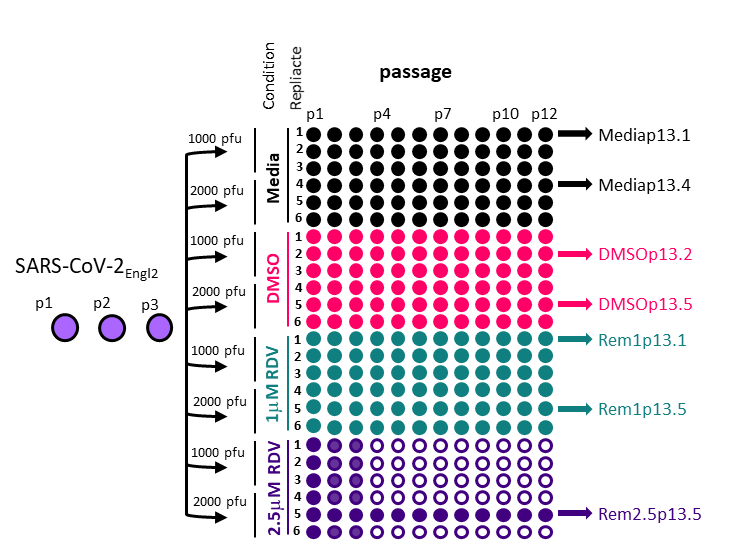

**Fig. S2. Schematic of the experimental layout to select for RDV resistant viruses.** The passage number of the input virus SARS-CoV-2_Engl2_ is given. P1 is the stock supplied by Public Health England, it was passaged twice in VeroE6 prior to selection in RDV. SARS-CoV-2_Engl2_ p3 is the input stock that was sequenced and used for analysis. Each condition is given a different colour and the amount of virus used at the start of the experiment is indicated. Populations that failed to amplify (no CPE or/and virus titre) are indicated by the empty circles. All lineages sequenced are indicated.

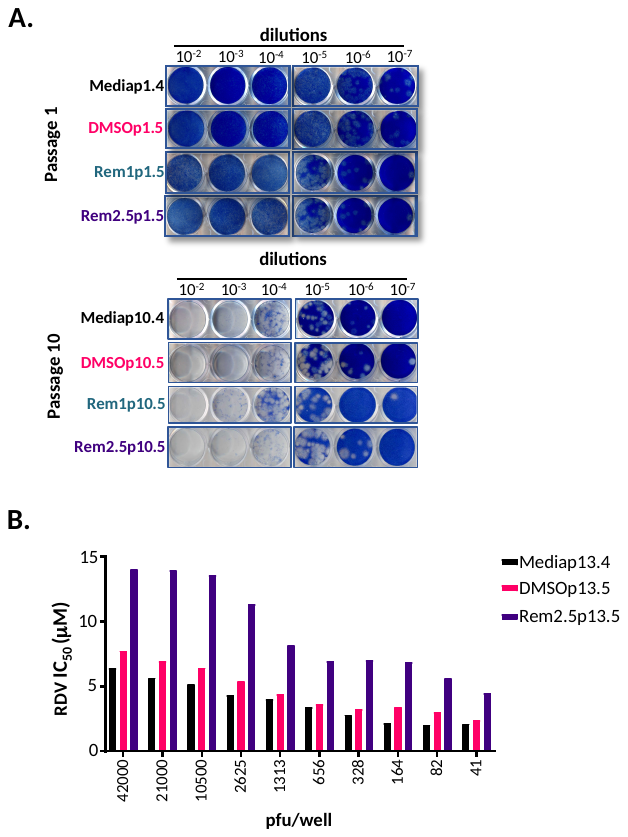

**Fig. S3. Adaptation of SARS-CoV-2 to VeroE6 and selection of decreased RDV sensitivity. A.** Coomassie blue stained SARS-CoV-2 plaque assays in VeroE6. Change in plaque morphology from passage 1 and passage 10. A subset of populations from the 4 different conditions were compared. **B.** Bar graph of RDV IC_50_ for a RDV adapted population and two control populations over a range of virus inputs.

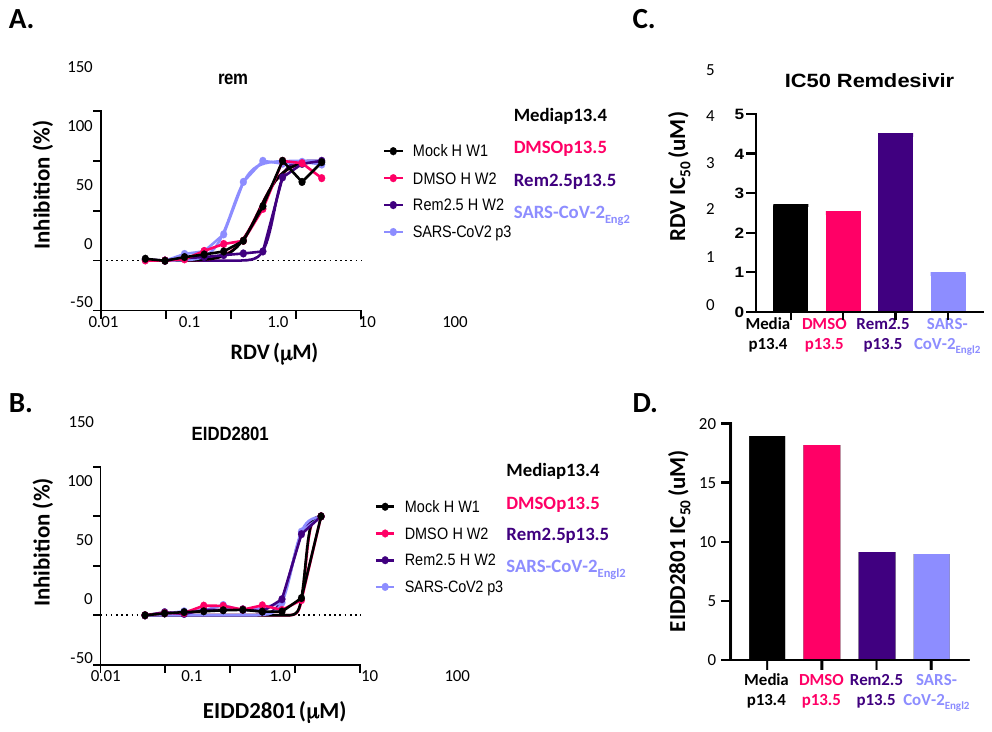

**Fig. S4.** **Antiviral activity of RDV and EIDD2801against RDV adapted virus populations**. RDV IC_50_ for the adapted population REM2.5p13.5 compared to the input virus SARS-CoV-2_Engl2_ as well as controls for adaption; DMSOp13.4 and Mediap13.4 in VeroE6-ACE2. **A.** RDV dose dependency curve. **B.** EIDD2801 dose dependency curve. **C.** Bar graph of RDV IC_50_ required to protect the monolayer. **D.** Bar graph of EIDD2801 IC_50_ required to protect the monolayer.

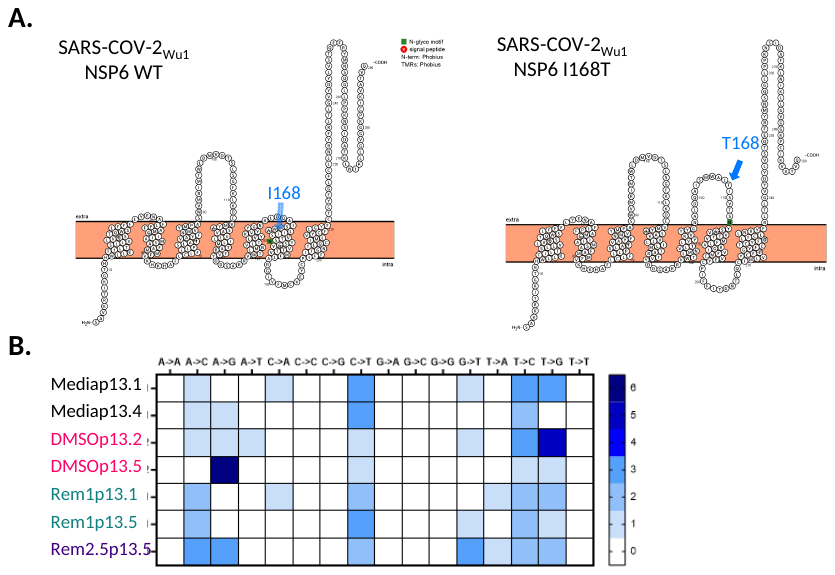

**Fig. S5.** **Sequence analysis of continually RDV selected SARS-CoV-2_Engl2_. A.** Heatmap of the distribution of transversion and transition depending on nucleotide. **B.** Protter prediction transmembrane organization of NSP6 wt (left) and NSP6 I168T (right) mutation. Position of I168 and I168T are indicated by an arrow. Extra-, intra-cellular and membrane is indicated, and putative glycosylation site is shown in green.

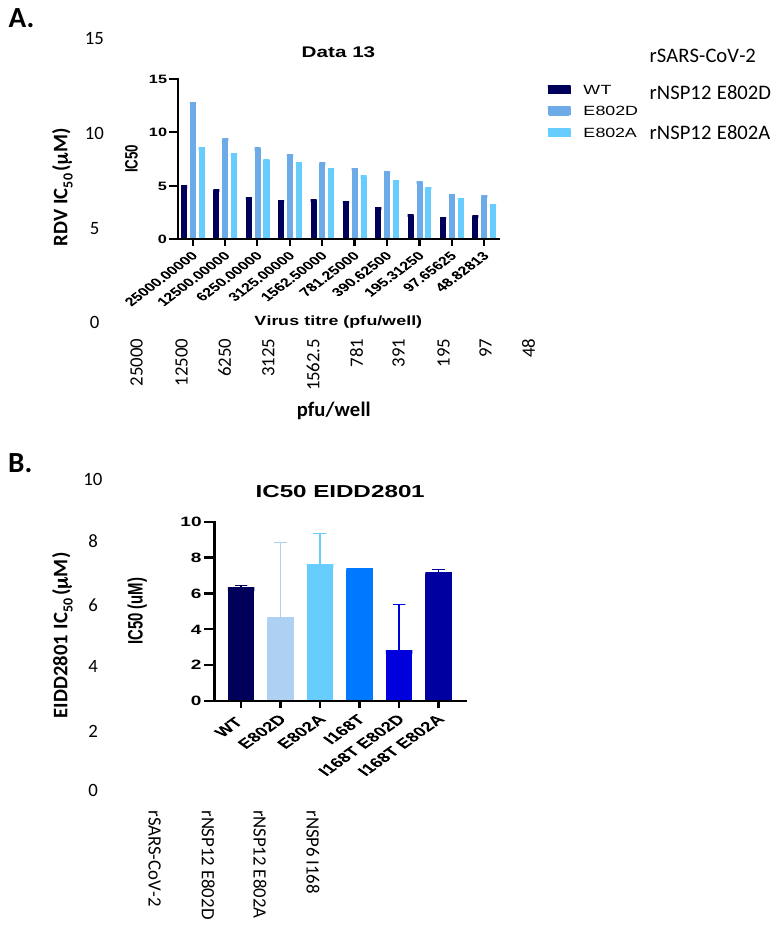

**Fig. S6.** **Antiviral activity of RDV and EIDD2801against NSP6 and NSP12 mutant viruses.** All viruses have a SARS-CoV-2_Wu1_ backbone with specific point mutations as indicated. Assay undertaken in VeroE6-ACE2-TMPRSS2. **A.** Bar graph of RDV IC_50_ values for rNSP12E902A, rNSP12E802D and rSARS-CoV-2 over a range of virus inputs. **B.** Bar graph of EIDD2801 IC_50_ for each rescued virus.

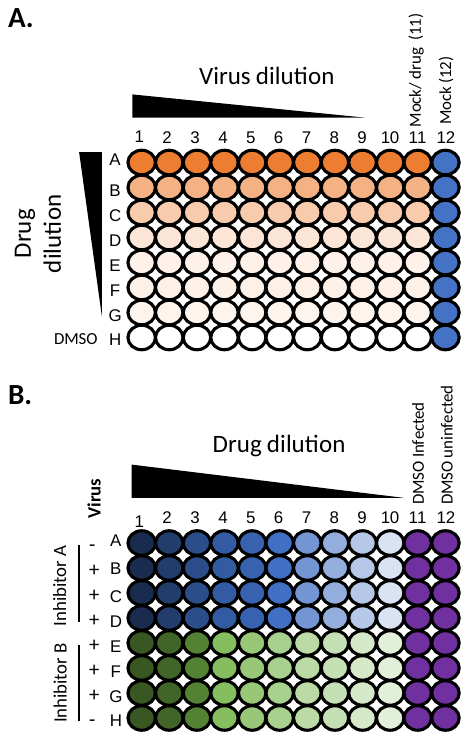

**Fig. S7.** **Schematic layout of 96-well plate clearance assay for drug screen**. All layouts have controls for inhibitor toxicity, virus clearance and cell monolayer integrity (DMSO only). **A.** Mixed array layout with range of inhibitor dilutions vs a range of virus dilutions. **B.** Mixed plate layout inhibitor was diluted and a set pfu/well. Amount of virus added causes complete well clearance by 72h pi. The mixed plate layout can also be modified to dilute inhibitor down the rows rather than across the columns.

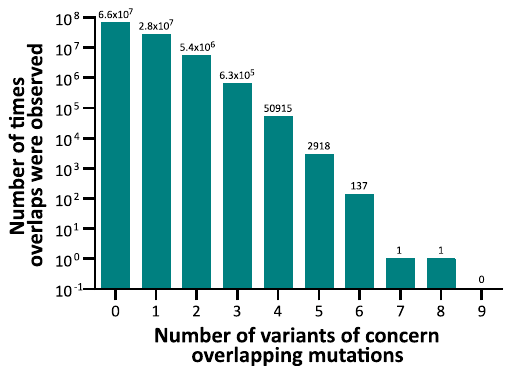

**Fig. S8.** **Distribution of convergence.** The null distribution of overlapping *in vitro* mutations hitting the same spike codon positions as in the SARS-CoV-2 variants of concern. 21 *in vitro* mutations and 20 variant of concern mutations spread across 1271 codons. 100 million simulations were run.

Table S1. RDV IC_50_ and fold-change in VeroE6-ACE2-TMPRSS2. All viruses have a SARS-CoV-2_Wu1_ backbone with specific point mutations as indicated. Average values from two independently rescued virus stocks, except for E802A. Experiment undertaken twice in triplicate by two independent operators.

| **Virus** | **IC_50_ (μM)** | **Fold-Change** |
| --- | --- | --- |
| rSARS-CoV-2 | 2.298 | - |
| rNSP12E802D | 5.676 | 2.47 |
| rNSP12E802A | 4.813 | 2.097 |
| rNSP6I168T | 2.417 | 1.05 |
| rNSP6I168T+NSP12E802D | 3.096 | 1.692 |
| rNSP6I168T+NSP12E802A | 3.728 | 1.347 |

Table S2. Average EIDD2801 IC_50_ and fold-change in VeroE6_ACE2_TMPRSS2. Average values from two independently rescued virus stocks, except for E802A Stock 1. Experiment undertaken twice in triplicate by two independent operators.

| **Virus** | **IC_50_ (μM)** | **Fold-Change** |
| --- | --- | --- |
| rSARS-CoV-2 | 6.348 | - |
| rNSP12E802D | 4.626 | 0.72 |
| rNSP12E802A | 7.388 | 1.19 |
| rNSP6I168T | 7.393 | 1.16 |
| rNSP6I168T+NSP12E802D | 2.816 | 0.44 |
| rNSP6I168T+NSP12E802A | 7.183 | 1.13 |

Table S3. dN/dS ratio

|  | **AllORFSeqs** | | | **ORF1ab** | | | **Spike** | | |
| --- | --- | --- | --- | --- | --- | --- | --- | --- | --- |
| **Sample** | **Syn** | **NonSyn** | **dN/dS** | **Syn** | **NonSyn** | **dN/dS** | **Syn** | **NonSyn** | **dN/dS** |
| Mediap13.1 | 4 | 7 | 0.5 | 2 | 2 | 0.25 | 0 | 4 | Infinity |
| Mediap13.4 | 0 | 6 | Infinity | 0 | 3 | Infinity | 0 | 3 | Infinity |
| DMSOp13.2 | 1 | 11 | 2.5 | 1 | 5 | 1.5 | 0 | 2 | Infinity |
| DMS0p13.5 | 2 | 6 | 1 | 0 | 2 | Infinity | 2 | 4 | 0.54 |
| Rem1p13.1 | 2 | 7 | 1 | 1 | 1 | 0.5 | 1 | 6 | 1.66 |
| Remp1p13.5 | 0 | 8 | Infinity | 0 | 6 | Infinity | 0 | 2 | Infinity |
| Rem2.5p13.5 | 2 | 13 | 2 | 0 | 5 | Infinity | 1 | 5 | 1.41 |

Data S1. Excel sheet summarizing of mutations across different continually passaged virus populations in comparison to SARS-CoV-2 _Wuhan-1_ (MN908947) and SARS-CoV-2_Engl2._

Data S2 Conservation of NSP12 E802 and NSP6 I168 in coronaviruses. Excel file sheet 1 frequency of NSP12 E802 and sheet 2 frequency of NSP6 I168. T
